# The EDC4-XRN1 axis controls P-body dynamics to link mRNA decapping with decay

**DOI:** 10.1101/2023.03.06.531261

**Authors:** William R. Brothers, Farah Ali, Sam Kajjo, Marc R. Fabian

**Affiliations:** Lady Davis Institute for Medical Research, Jewish General Hospital, Montreal, Quebec, Canada; Department of Biochemistry, McGill University, Montreal, Quebec, Canada; Department of Oncology, McGill University, Montreal, Quebec, Canada

## Abstract

Deadenylation-dependent mRNA decapping and decay is the major cytoplasmic mRNA turnover pathway in eukaryotes. Many mRNA decapping and decay factors associate with each other via protein-protein interaction motifs. For example, the decapping enzyme DCP2 and the 5’-3’ exoribonuclease XRN1 interact with enhancer of mRNA decapping protein 4 (EDC4), a large scaffold that has been reported to stimulate mRNA decapping. mRNA decapping and decay factors are also found in processing bodies (P-bodies), evolutionarily conserved ribonucleoprotein (RNP) granules that are often enriched with mRNAs targeted for decay, such as microRNA (miRNA)-targeted mRNAs, yet paradoxically are not required for mRNA decay to occur. In this study, we show that disrupting the interaction between XRN1 and EDC4 or altering their stoichiometry leads to an inhibition of mRNA decapping, with miRNA-targeted mRNAs being stabilized in a translationally repressed state. Importantly, we demonstrate that this concomitantly leads to larger P-bodies that are directly responsible for preventing mRNA decapping under these conditions. Finally, we demonstrate that P-bodies act to support cell viability and prevent stress granule formation under conditions when XRN1 is limiting. Taken together, these data demonstrate that the interaction between XRN1 and EDC4 regulates P-body dynamics to properly coordinate mRNA decapping with 5’-3’ decay in human cells.

**HIGHLIGHTS:**

- XRN1-EDC4 interaction couples mRNA decapping with mRNA decay.
- Disrupting XRN1-EDC4 contact generates larger P-bodies that, in turn, inhibit decapping.
- P-bodies support cellular fitness in the absence of XRN1.

## INTRODUCTION

Translation and mRNA stability are tightly controlled to post-transcriptionally regulate gene expression. In eukaryotes, the majority of cytoplasmic mRNA decay occurs through a deadenylation-dependent program, which involves the removal of the poly(A) tail from the 3’ end of the mRNA (Jonas & Izaurralde, 2015). This is accomplished by deadenylase machineries, including the PAN2-PAN3 and CCR4-NOT deadenylase complexes. Multiple mRNA repression programs directly recruit the CCR4-NOT complex to the 3’ untranslated regions (UTRs) of targeted mRNAs through various 3’UTR binding proteins. For example, microRNA (miRNA) target sites associate with the miRNA induced silencing complex (miRISC), which recruits the CCR4-NOT complex to initiate mRNA deadenylation (Braun *et al*, 2011; Collart & Panasenko, 2012; Fabian *et al*, 2010; Jonas & Izaurralde, 2015). Following deadenylation, the mRNA decapping complex is recruited to the 5’-terminus of the mRNA (Yamashita *et al*, 2005). This includes the DCP2 decapping enzyme, the decapping cofactor DCP1, and enhancer of decapping proteins such as EDC3 and EDC4 (Chang *et al*, 2014; Chang *et al*, 2019; Vidya & Duchaine, 2022), which function to hydrolyze the N^7^-methylguanosine 5’-cap structure. Following hydrolysis of the 5’-cap, the mRNA is vulnerable to degradation by the 5’-3’ exonuclease XRN1, which physically associates with the mRNA decapping complex (Braun *et al*, 2012; Chang *et al*., 2014; Topisirovic *et al*, 2011).

Deadenylation, decapping, and decay factors interact with each other via a complex network of multivalent protein-protein interactions, which can act to support mRNA turnover (Jonas & Izaurralde, 2013). In addition, proteins with described roles in mRNA decay and translational repression can be spatially organized in cytoplasmic ribonucleoprotein (RNP) granules known as processing bodies (P-bodies). P-bodies are dynamic structures that form through complex networks of protein-protein, protein-RNA, and RNA-RNA interactions between RNA-binding proteins, and their cognate mRNAs (Standart & Weil, 2018). Indeed, several mRNA decapping and decay factors act as core nucleating proteins critical for visible P-body formation. These include EDC4, the eIF4E-binding protein 4E-T, DCP1, or Like Sm14 (LSM14) (Franks & Lykke-Andersen, 2008; Jonas & Izaurralde, 2013; Parker & Sheth, 2007). For example, LSM14 directly interacts with EDC4, 4E-T, and DDX6, and disrupting any of these interactions prevents visible P-body assembly (Brandmann *et al*, 2018).

Despite being highly concentrated in mRNA decay factors, P-bodies are enigmatic in that they are not required for mRNA decay to occur (Eulalio *et al*, 2007b). Recently, several groups have made correlative observations that suggest there may be a link between P-bodies and the negative regulation of mRNA decapping and 5’ to 3’ decay (Hubstenberger *et al*, 2017; Na *et al*, 2020; Tibble *et al*, 2021). Moreover, mRNA decapping subunit stoichiometry has been linked to P-body assembly. Interestingly, though, while mRNA decapping is disrupted by depleting cells of any of the mRNA decapping complex subunits, individual subunit depletion can lead to disparate effects on P-body formation. For instance, DCP1 and EDC4 are both core P-body proteins, where each of their depletion leads to the loss of P-bodies (Aizer *et al*, 2013; Eulalio *et al*, 2007a; Eulalio *et al*., 2007b; Luo *et al*, 2018). Conversely, depleting cells of DCP2 or XRN1 results in an increase in P-body size in *Drosophila* and yeast cells (Eulalio *et al*., 2007b; Sheth & Parker, 2003). However, despite these observations, direct mechanistic evidence to support a role for P-bodies in regulating mRNA decapping and decay is lacking. Moreover, precisely what regulates the coordination between mRNA decapping and 5’ to 3’ decay in human cells remains unknown.

Here, we set out to elucidate how changes in the levels of mRNA decapping complex subunits regulates miRNA-mediated mRNA decay. Interestingly, we observe that altering the stoichiometry of EDC4 and XRN1 results a shift in the mode of miRNA-silencing from mRNA decay to translational repression. Specifically, we show that upregulating EDC4 levels, depleting XRN1, or disrupting the ability of EDC4 to interact with XRN1 generates larger P-bodies. We go on to show that the enlarged P-bodies directly stabilize miRNA-targeted mRNAs by inhibiting mRNA decapping under these conditions. Finally, we demonstrate that P-bodies can provide a fitness advantage in the absence of XRN1, as disrupting P-body formation in XRN1 knockout cells leads to stress granule formation and reduces cell viability. Taken together, our data support a model where the EDC4-XRN1 interaction regulates mRNA decapping activity through P-body formation, which feeds back on cell viability when XRN1 is limiting.

## RESULTS

### Elevated EDC4 levels shifts the mode of miRNA-mediated silencing from mRNA decay to translational repression by remodeling P-bodies

The depletion of EDC4 has been linked to a loss of mRNA decapping activity and decay (Erickson *et al*, 2015). Therefore, we hypothesized that increasing cellular EDC4 levels might enhance the decay of a miRNA-targeted reporter. To test this, EDC4-overexpressing HeLa cells were transfected with plasmids encoding a *Renilla* luciferase (RL) reporter mRNA with six *let-7* miRNA target sites (RL-6xB) or six mutated let-7 sites (RL-6xBMUT) in its 3’UTR (**Figure 1A and B**) (Fabian *et al*, 2009). 48 hours post-transfection, cells were lysed and RL-6xB silencing was measured using luciferase assays, with RL-6xBMUT serving as a control. In addition, RL-6xB and RL-6xBMUT mRNA stability was assessed by Actinomycin D (Act-D) treatment to inhibit transcription, isolation of total RNA at multiple time points, and measurement of mRNA levels by reverse transcription-quantitative PCR (RT-qPCR). Overexpression of EDC4 did not significantly alter the extent of RL-6xB repression as determined by luciferase assays (**Figure 1C**). Surprisingly, though, while RL-6xB mRNA rapidly degraded in cells expressing only endogenous EDC4, RL-6xB mRNA was stabilized in EDC4-overexpressing cells with a decay rate similar to that of the RL-6xBMUT control mRNA (**Figure 1D**).

**Figure 1.**
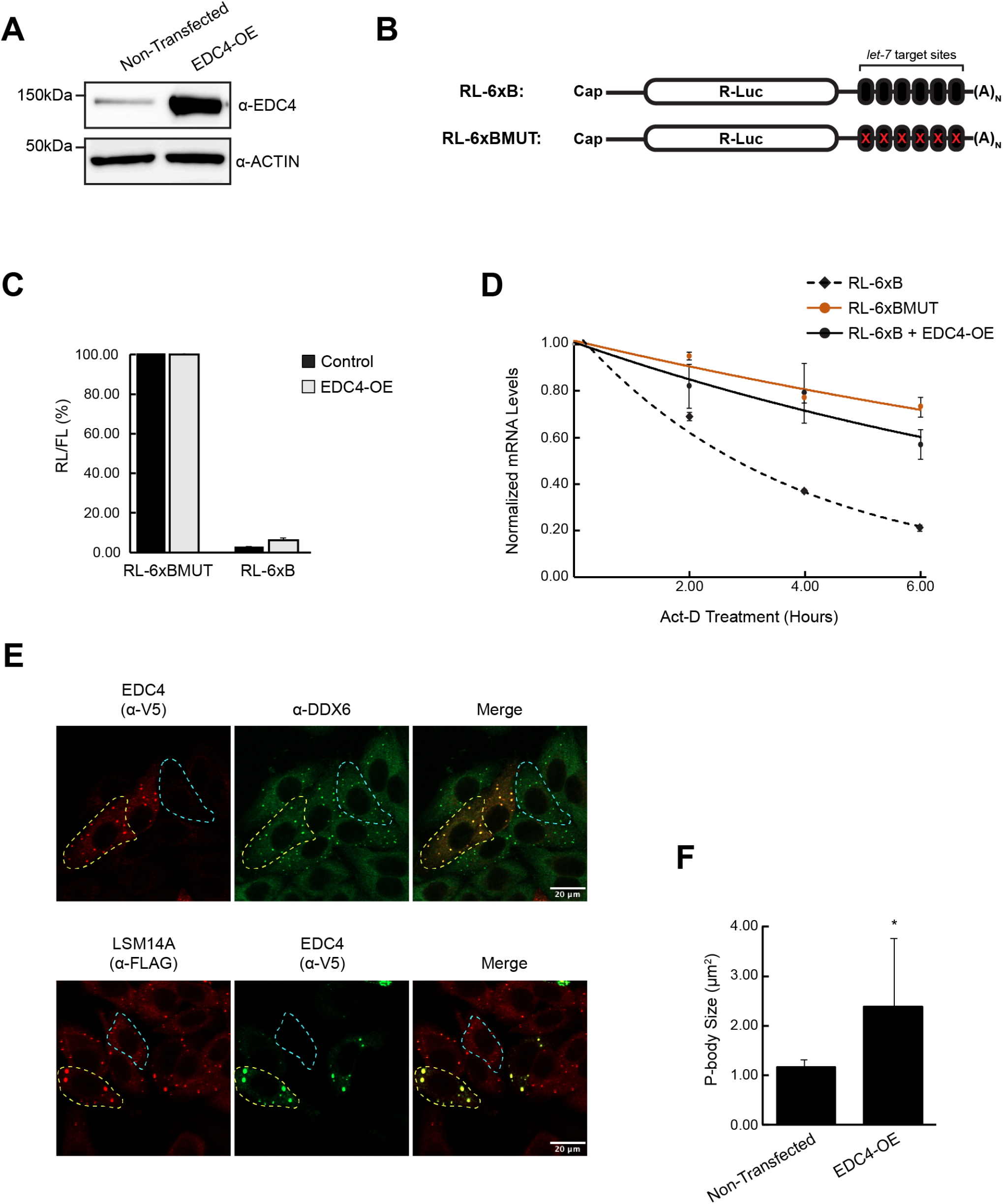
EDC4 levels enhance P-body condensation and stabilize miRNA-targeted mRNAs. (**A**) Western blot validation of EDC4 overexpression. (**B**) Schematic diagrams of *Renilla* luciferase reporter mRNAs with six *let-7* (RL-6xB) or mutated *let-7* (RL-6xBMUT) target sites. (**C**) Luciferase assays performed on HeLa cells expressing the indicated reporters with or without EDC4-overexpression. Luciferase values are expressed as a RL/FL ratio with the RL/FL ratio of RL-6xBMUT expressing cells normalized to 100%. Error bars represent the SEM of three biological replicates. (**D**) mRNA decay assay assessing the decay rates of cells expressing the indicated reporters and proteins. RNA was isolated from cells after treatment with Actinomycin D (Act-D) to halt transcription. mRNA levels were measured by RT-qPCR with RL-6xB levels being normalized to *GAPDH* levels. Normalized RL-6xB levels at zero hours of Act-D treatment were set to 1.0. Error bars represent the SEM of three biological replicates. (**E**) Images generated through confocal microscopy of HeLa cells transfected with plasmids encoding EDC4-V5. Immunofluorescent staining was performed using ⍺-V5 and antibodies to detect the P-body markers DDX6 and LSM14A. Images were processed, and channels merged using Fiji. Yellow outlines highlight cells that have been transfected with the indicated proteins, in contrast to non-transfected cells (blue outlines). (**F**) Quantification of P-body sizes in cells transfected with EDC4-V5 compared to non-transfected controls in the same samples. Measurements were performed on three separate biological replicates using Fiji. Error bars represent the SEM and statistical significance was established using a two-tailed student’s T-test (* = P < 0.05).

Previously, we showed that EDC4 overexpression in HeLa cells leads to an increase in P-body size (**Figures 1E and F**) (Brothers *et al*, 2022; Brothers *et al*, 2020). Recent single-molecule imaging studies point to P-bodies serving as “waystations” for the persistent translational repression of mRNAs targeted by the miRISC (Cialek *et al*, 2022). Moreover, several lines of evidence suggest that mRNAs within P-bodies are not actively undergoing decay (Arribere *et al*, 2011; Di Stefano *et al*, 2019; Eulalio *et al*., 2007b; Hubstenberger *et al*., 2017). We therefore wanted to determine if EDC4-remodeled P-bodies play a role in stabilizing the RL-6xB mRNA. To this end, we targeted the core P-body protein LSM14A using CRISPR-Cas9 to generate knock-out LSM14A HeLa cells (LSM14A^KO^) and then stably rescued LSM14A expression with wild-type FLAG-tagged LSM14A (F-LSM14A^WT^) or LSM14A mutants that do not interact with EDC4 (F-LSM14A^ΔFFD^) or DDX6 (F-LSM14A^ΔTFG^) (**Figures 2A and B**) (Brandmann *et al*., 2018). Importantly, these LSM14A mutants do not support visible P-bodies, even when EDC4 is overexpressed (**Figure 2C**) (Brothers *et al*., 2022). Cells were transfected with RL-6xB and RL-6xBMUT reporter constructs alone or with a plasmid encoding V5-tagged EDC4. We found that RL-6xB was repressed equally well in F-LSM14A^WT^ or F-LSM14A^ΔFFD^ cells as assessed by luciferase assays, regardless of EDC4 overexpression (**Figure 2D**). However, while RL-6xB mRNA was stabilized in F-LSM14A^WT^ cells that expressed high levels of EDC4, it rapidly decayed in F-LSM14A^ΔFFD^ and F-LSM14A^ΔTFG^ cells that lack P-bodies (**Figures 2E**). We also examined the decay rates of *MCL1* and *BCL2*, two endogenous mRNAs that rapidly decay and are reported to be enriched in P-bodies (Cui & Placzek, 2018; Hubstenberger *et al*., 2017; Ishimaru *et al*, 2010). Like RL-6xB, EDC4 overexpression stabilized both *MCL1* and *BCL2* mRNAs in F-LSM14A^WT^ cells but had no effect in LSM14A^ΔFFD^ cells with both transcripts decaying in a rapid manner (**Figure 2F and S1A-B**). Using an alternative approach, we disrupted P-bodies using NBDY, a microprotein that interacts with EDC4 and DCP1A and promotes P-body dissociation (**Figures 3A and B**) (Brothers *et al*., 2022; D’Lima *et al*, 2017; Na *et al*, 2021; Na *et al*., 2020). Expressing NBDY in EDC4-overexpressing cells also restored the decay of exogenously introduced RL-6xB, or endogenous *MCL1* and *BCL2* mRNAs to rates similar to those observed in control cells (**Figures 3C-E**). Taken together, our results indicate that upregulating EDC4 remodels P-body condensates, which in turn is responsible for shifting the mode of miRNA-mediated silencing from mRNA decay to translational repression.

**Figure 2.**
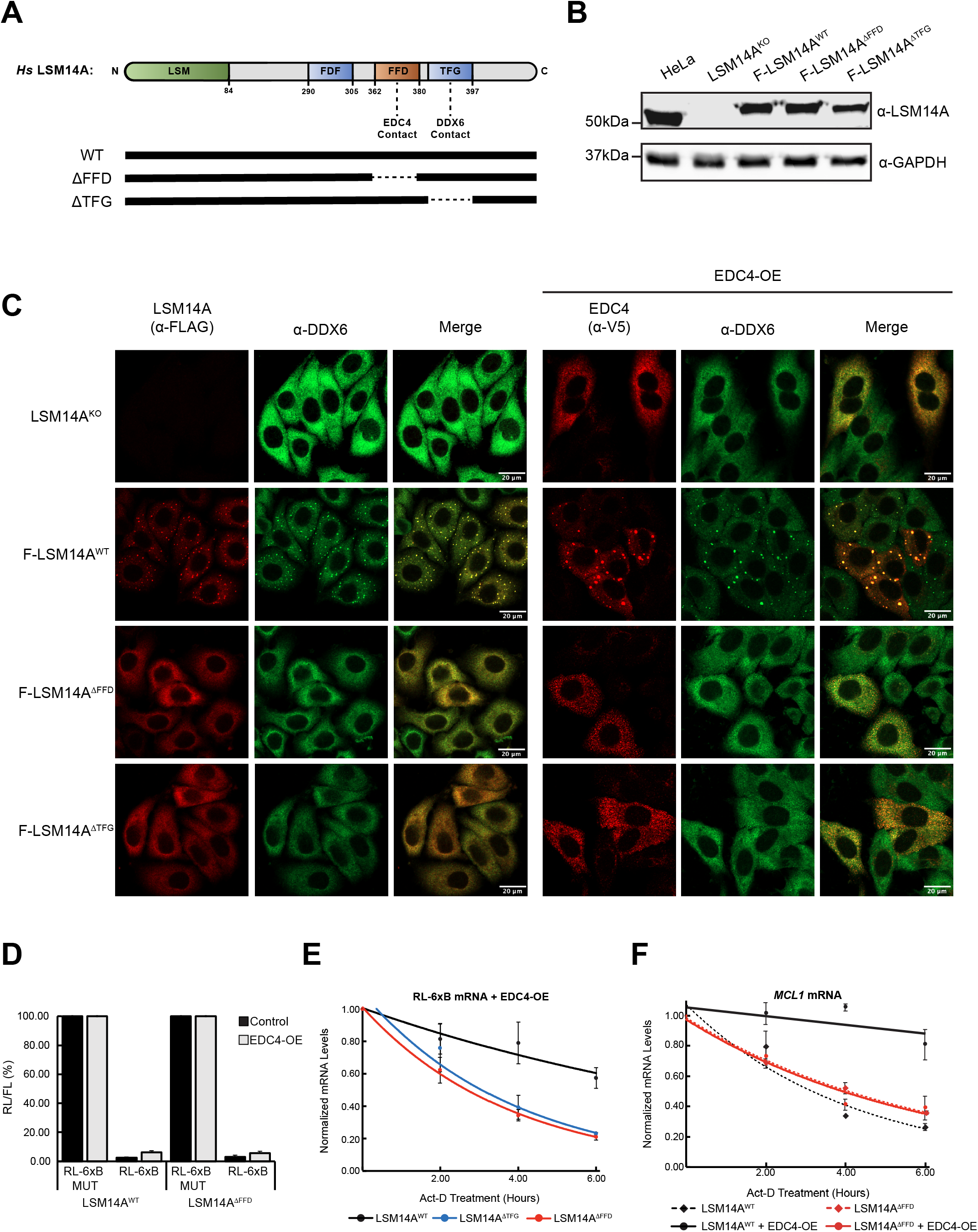
P-bodies bias miRNA-silencing from target mRNA decay to translational repression. (**A**) Schematic diagram of human LSM14A, along with diagrams representing deletion mutants that disrupt P-body formation. (**B**) Western blotting of WT or LSM14^KO^ HeLa cells stably rescued with FLAG-tagged (F-)LSM14A proteins. (**C**) Images generated through confocal microscopy of LSM14A^KO^ HeLa cells stably expressing F-LSM14A constructs under endogenous EDC4 expression or EDC4-overexpression. Immunofluorescent staining was performed using ⍺-FLAG under endogenous EDC4 expression levels or ⍺-V5 under EDC4 overexpression with ⍺-DDX6. Images were processed, and channels merged using Fiji. (**D**) Luciferase assays performed on LSM14A^WT^ or LSM14A^ΔFFD^ cells expressing the indicated reporters with or without EDC4-overexpression (EDC4-OE). Luciferase values are expressed as a RL/FL ratio with the RL/FL ratio of RL-6xBMUT expressing cells normalized to 100%. Error bars represent the SEM of three biological replicates. (**E-F**) mRNA decay assays in the indicated F-LSM14 cell lines assessing the decay rates of miRNA-targeted mRNAs with or without EDC4-OE. RNA was isolated from cells after treatment with Actinomycin D (Act-D) to halt transcription. mRNA levels were measured by RT-qPCR with RL-6xB (**E**) or *MCL1* (**F**) levels being normalized to *GAPDH* levels. Normalized RL-6xB (**E**) or *MCL1* (**F**) levels at zero-hours of Act-D treatment were set to 1.0. Error bars represent the SEM of three biological replicates.

**Figure 3.**
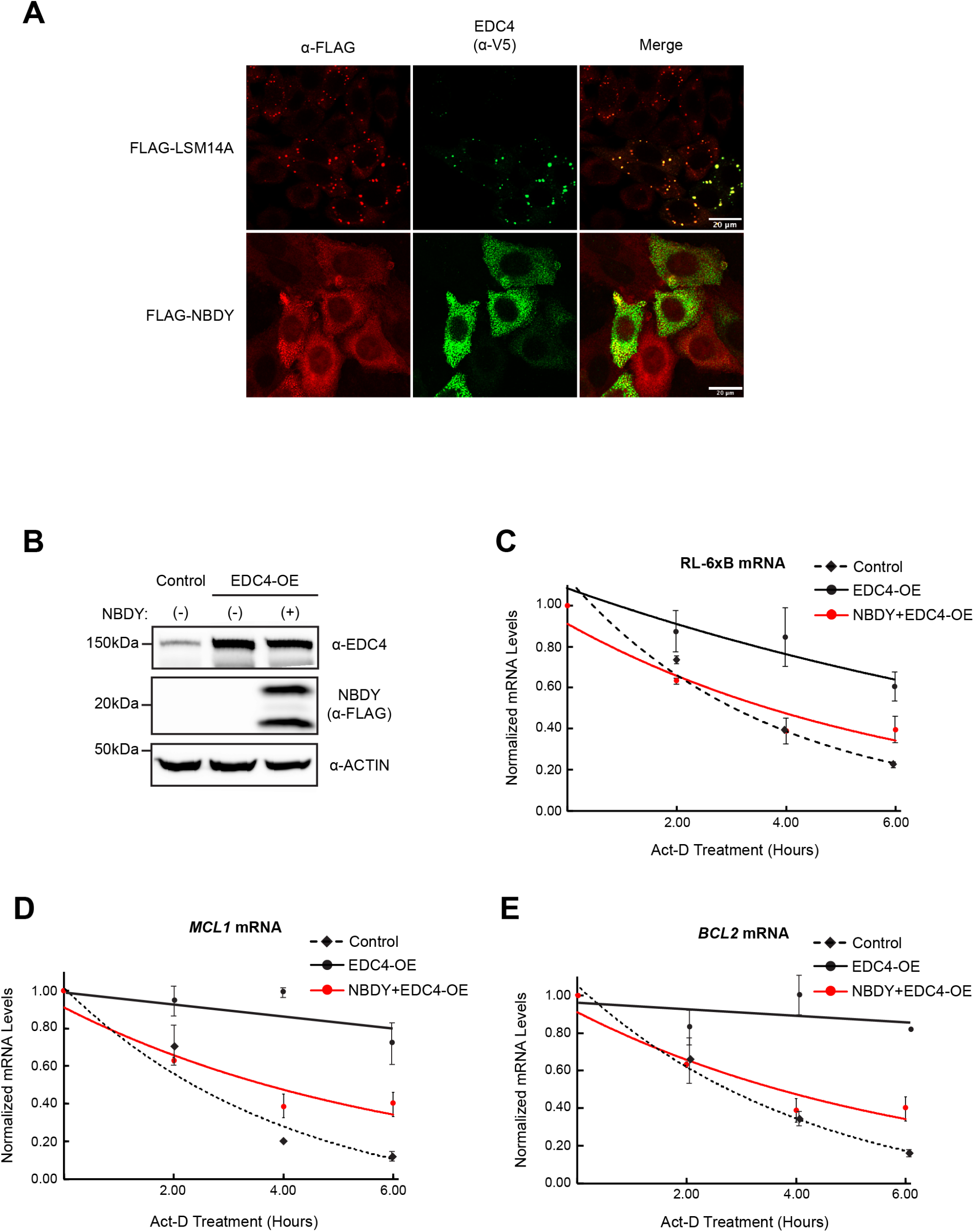
NBDY negates the effects of EDC4-overexpression (EDC4-OE) on miRNA-mediated decay rates. (**A**) Images generated through confocal microscopy of HeLa cells transfected with plasmids encoding EDC4-V5 and FLAG-NBDY. Immunofluorescent staining was performed using ⍺-FLAG and ⍺-V5 antibodies. Images were processed, and channels merged using Fiji. (**B**) Western blots of HeLa cell extracts expressing the indicated proteins. (**C-E**) mRNA decay assay assessing the decay rates of miRNA-targeted mRNAs with or without EDC4-OE. RNA was isolated from cells after treatment with Actinomycin D (Act-D) to halt transcription. mRNA levels were measured by RT-qPCR with RL-6xB (**C**), *MCL1* (**D**), or *BCL2* (**E**) levels being normalized to *GAPDH* levels. Normalized RL-6xB (**C**), *MCL1* (**D**), or *BCL2* (**E**) levels at zero-hours of Act-D treatment were set to 1.0. Error bars represent the SEM of three biological replicates.

### EDC4 overexpression enriches stabilized miRNA-targeted mRNAs within P-bodies

Both exogenously introduced reporters and endogenous mRNAs targeted by miRNAs have been reported to localize to P-bodies (Hubstenberger *et al*., 2017; Liu *et al*, 2005). As our data indicate that overexpressing EDC4 stabilizes a miRNA-targeted mRNA by enhancing P-body formation, we next wanted to determine if miRNA-targeted mRNAs are differentially localized to P-bodies under these conditions. To assess this, we transfected F-LSM14A^WT^ HeLa cells with plasmid encoding RL-6xB alone or with plasmids that overexpress V5-tagged EDC4 and performed single-molecule fluorescent *in situ* hybridization (smFISH) using Cy3-labeled probes against the RL-6xB reporter mRNA. To establish whether the smFISH signal co-localizes to *bona fide* P-bodies, we performed simultaneous immunofluorescent staining against DDX6 and either LSM14A (⍺-FLAG) when EDC4 was not overexpressed or V5-EDC4 (⍺-V5) in EDC4-overexpressing cells (**Figure 4A**). As overexpressing EDC4 only stabilized RL-6xB mRNA in cells that can form visible P-bodies, we also treated transfected cells with Act-D for 4 hours to investigate whether RL-6xB mRNAs are differentially stabilized within EDC4-remodelled P-bodies (**Figure 4B**).

**Figure 4.**
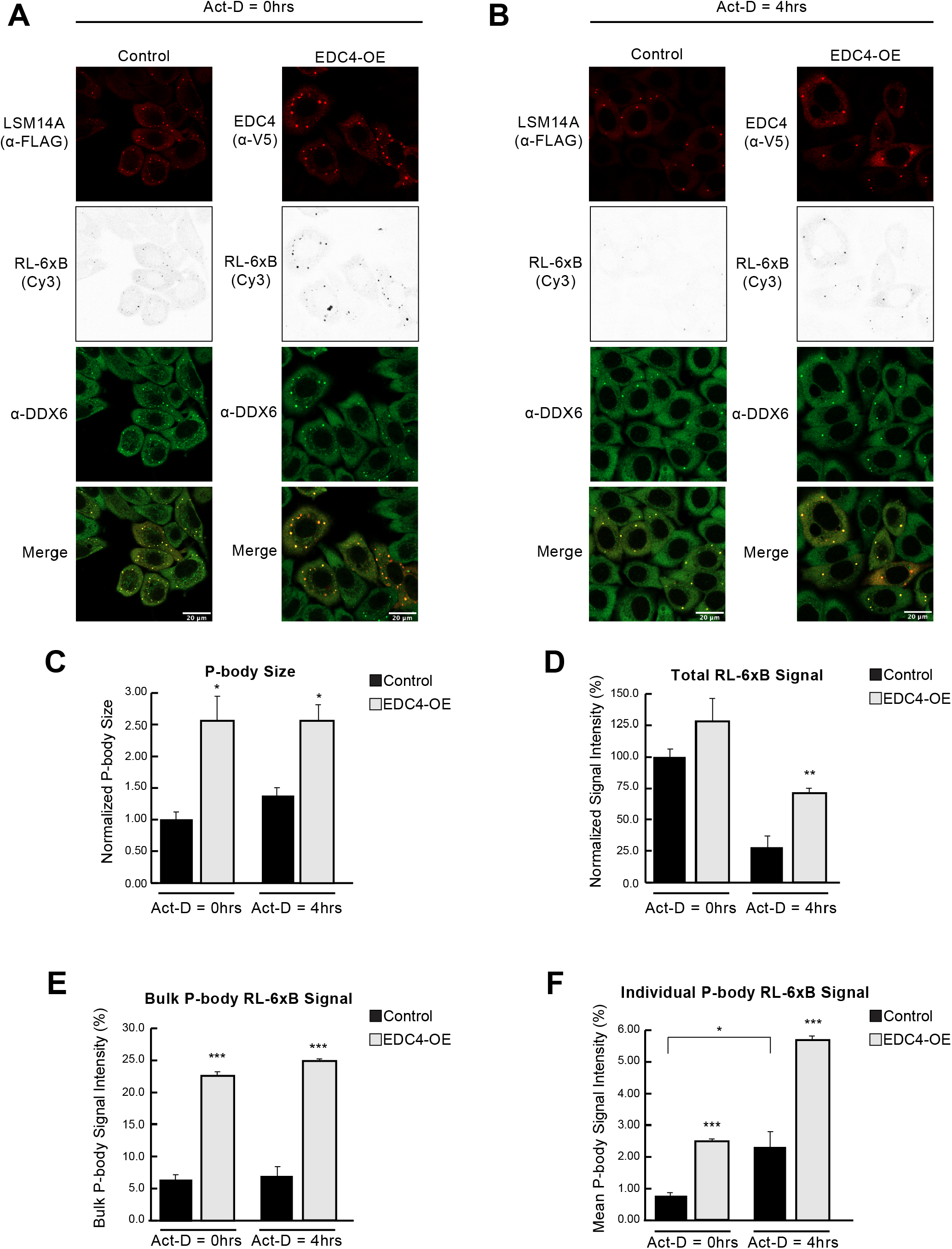
EDC4-overexpression differentially recruits miRNA-targeted mRNAs to P-bodies and protects them from decay. (**A-B**) Images generated through confocal microscopy of F-LSM14A^WT^ HeLa cells transfected with plasmids encoding EDC4-V5 after zero (**A**) or four hours (**B**) of Actinomycin D (Act-D) treatment. Single-molecule fluorescent *in situ* hybridization (smFISH) using Cy3-labeled probes against RL-6xB with simultaneous immunofluorescent staining was performed using ⍺-DDX6 with ⍺-FLAG or ⍺-V5 antibodies. Images were processed, analyzed, and channels merged using Fiji. (**C**) Quantification of P-body sizes from experiments depicted in (**A-B**) with the mean P-body size of control cells with zero hours of Act-D treatment set to 100%. Measurements were performed on three separate biological replicates using Fiji. Error bars represent the SEM and statistical significance was established using a two-tailed student’s T-test relative to control conditions (* = P < 0.05). (**D**) Quantification of total whole cell smFISH signal from the RL-6xB Cy3 channel from experiments depicted in (**A-B**) with the mean smFISH signal of control cells with no Act-D treatment set to 100%. Measurements were performed on three separate biological replicates using Fiji. Error bars represent the SEM and statistical significance was established using a two-tailed student’s T-test relative to control conditions (** = P < 0.01). (**E**) Quantification of the additive smFISH signal from all P-bodies as a fraction of the condition-matched total smFISH signal from experiments depicted in (**A-B**). Measurements were performed on three separate biological replicates with ten cells per replicate using Fiji. Error bars represent the SEM and statistical significance was established using a two-tailed student’s T-test relative to control conditions (*** = P < 0.001). (**F**) Quantification of the additive smFISH signal per individual P-body as a fraction of the condition-matched total smFISH signal from experiments depicted in (**A-B**). Measurements were performed on three separate biological replicates with ten cells per replicate using Fiji. Error bars represent the SEM and statistical significance was established using a two-tailed student’s T-test (* = P < 0.05; *** = P < 0.001).

P-bodies, as defined by the punctate signal overlap between the two protein channels, were approximately two-fold larger in EDC4-overexpression conditions (**Figure 4C**). We also stained for the stress granule marker G3BP1, which remained diffuse throughout the cytoplasm and did not overlap with RL-6xB smFISH signal (**Figure S1C**), suggesting that EDC4 overexpression does not lead to stress granule formation. Next, we quantified the overall smFISH signal from cells expressing RL-6xB. Consistent with our results from mRNA stability assays (**Figure 1F**), RL-6xB smFISH signal was significantly elevated in EDC4-overexpressing cells after 4 hours of Act-D treatment compared to cells expressing only endogenous EDC4 (**Figure 4D**). Additionally, we found significant enrichment of RL-6xB signal overlapping with P-bodies upon EDC4 overexpression relative to cells with endogenous EDC4 (**Figure 4E**). It is important to note that cells treated with Act-D exhibited fewer but larger P-bodies than untreated cells, as has been reported previously (Cougot *et al*, 2004). Therefore, we also measured the mean RL-6xB signal per individual P-body to normalize for differences in P-body numbers between conditions (**Figure 4F**). This analysis showed that Act-D treatment was associated with higher RL-6xB signal per P-body in both control and EDC4-overexpressing cells, indicating that mRNAs enriched in P-bodies are being preferentially protected from degradation. Together, these results suggest that RL-6xB mRNAs are enriched within P-bodies due to EDC4 overexpression, leading to their protection from decay.

### miRNA-targeted mRNAs stabilized in P-bodies are deadenylated but not decapped

miRNA-targeted mRNAs are generally destabilized via deadenylation, followed by decapping and subsequent 5’-3’ decay (Jonas & Izaurralde, 2015). Our data suggest that the decay of the *let-7* targeted RL-6xB mRNA is impaired when EDC4 enhances P-body condensates. We therefore set out to identify what step in the mRNA decay pathway is inhibited under these conditions. To assess the poly(A) tail status of RL-6xB mRNA in EDC4-overexpressing cells, we isolated total RNA at zero and six hours following Act-D treatment and performed enhanced poly(A) tailing (ePAT) assays (Jänicke *et al*, 2012) (**Figure 5A**). While RL-6xB mRNA displayed long heterogenous poly(A) tails at the zero hour timepoint, it maintained a short poly(A) tail after six hours of Act-D treatment, suggesting that these mRNAs are being actively targeted for deadenylation. This contrasts with *GAPDH* mRNAs, which are also stable but maintain heterogeneous poly(A) tail lengths at both zero and six hours of Act-D treatment. To determine the RL-6xB mRNA cap status, we incubated RNA isolated from cells treated with Act-D for six hours with Terminator nuclease, a 5’-3’ exonuclease that degrades uncapped RNAs with a 5’ monophosphate (Braun *et al*., 2012). Following Terminator treatment, total RNA was isolated and RNA levels were quantified by RT-qPCR with spike-in RNA for normalization (**Figure 5B**). The Terminator nuclease failed to efficiently degrade RL-6xB mRNA from samples collected after six hours of Act-D treatment, as these mRNAs were reduced to similar degrees as *GAPDH* mRNA. In contrast, the Terminator nuclease efficiently degraded 18S rRNA, which does not harbor a 5’-cap. However, RL-6xB mRNA levels were further reduced by Terminator nuclease when we first treated RNA *in vitro* with an mRNA decapping enzyme (Paquette *et al*, 2018) (**Figure S1D**). These results suggest that enhanced P-body formation prevents miRNA-targeted mRNAs from being decapped after they have been deadenylated.

**Figure 5.**
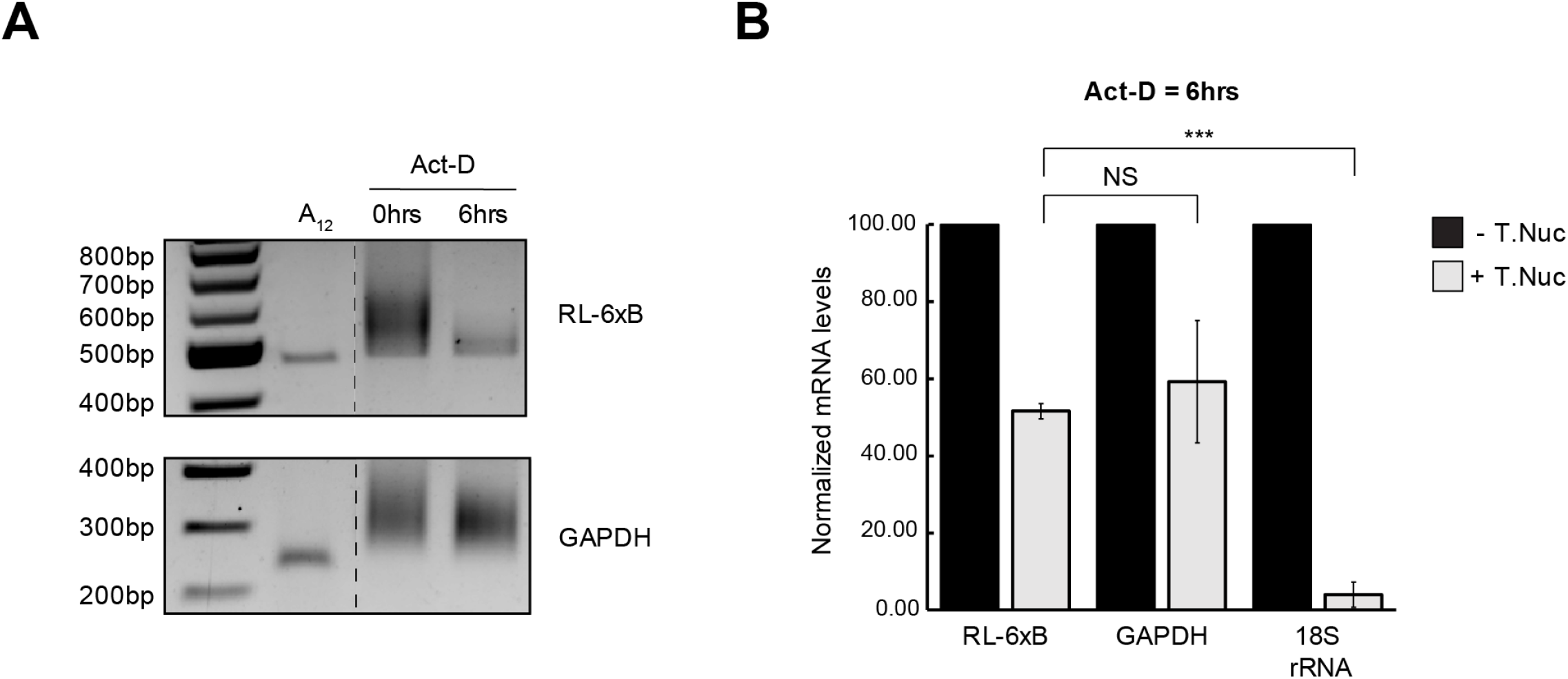
P-body condensates uncouple deadenylation from mRNA decapping. (**A**) Poly(A) tail analyses using an enhanced poly(A) tailing (ePAT) assay performed on RNA isolated from HeLa cells overexpressing EDC4 and treated with Actinomycin D (Act-D) for zero or six hours. A_12_ represents ePAT assays performed on mRNAs that have fixed length 12-adenine tails. (**B**) Cap status analysis of mRNAs collected from HeLa cells overexpressing EDC4 after 6-hours of Act-D treatment. Isolated RNA was incubated *in vitro* with terminator nuclease (T. Nuc) along with no-enzyme controls. Purified RNA underwent RT-qPCR analysis for the annotated genes normalized to an *in vitro* transcribed *firefly luciferase* mRNA spike-in control introduced at the RT step. Normalized mRNA levels of no-enzyme control samples were set to 100%, error bars represent the SEM of three biological replicates, and statistical significance was established using a two-tailed student’s T-test (NS = not significant; *** = P < 0.001).

### Depleting XRN1 inhibits mRNA decapping by enhancing P-body condensation

Our data suggest that enhancing P-body formation by upregulating EDC4 levels limits mRNA decapping in human cells. It has previously been shown that depleting XRN1 in yeast and *Drosophila* S2 cells leads to similarly enlarged P-bodies (Eulalio *et al*., 2007b; Sheth & Parker, 2003), a phenotype that we also observe in XRN1^KO^ HeLa cells compared to WT HeLa cells (**Figures 6A-C**). Moreover, in *Drosophila* S2 cells, depletion of XRN1 has been reported to inhibit mRNA decapping via an unknown mechanism (Braun *et al*., 2012). We therefore aimed to determine if loss of XRN1 impairs mRNA decapping in human cells, and if this effect is mediated by alterations in P-body condensation. To test this, we introduced the RL-6xB reporter into WT and XRN1^KO^ HeLa cells alone or with a plasmid encoding FLAG-tagged NBDY to disrupt visible P-bodies formation (**Figures 6D and E**). Cells were treated with Act-D for six hours to inhibit *de novo* transcription and total RNA was isolated to assess both mRNA stability and 5’ cap status, the latter by incubating isolated RNA with Terminator nuclease (**Figures 6F and G**). RL-6xB mRNA was significantly more stable XRN1^KO^ cells compared to WT cells, regardless of NBDY expression (**Figure 6F**). However, Terminator nuclease failed to degrade RL-6xB mRNA from XRN1^KO^ cells, but efficiently degraded RL-6xB mRNA from NBDY-expressing XRN1^KO^ cells (**Figure 6G**). The efficient degradation of ribosomal RNA by Terminator nuclease under both conditions suggests that the differences in RL-6xB mRNA degradation were not caused by changes in Terminator nuclease activity. Taken together, these results indicate that loss of XRN1 has a similar effect to overexpression of EDC4, resulting in impaired mRNA decapping by enhancing P-body formation in human cells.

**Figure 6.**
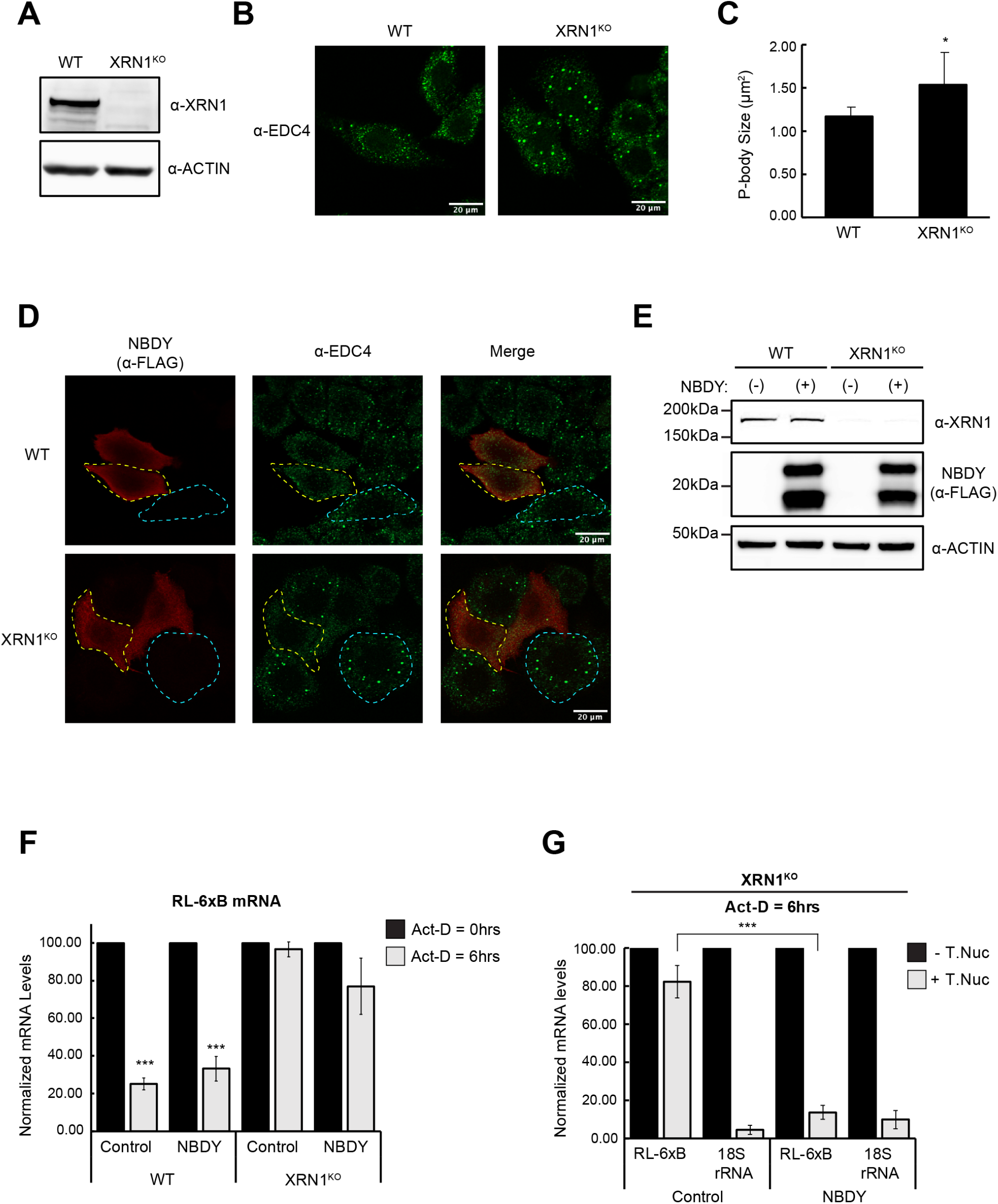
XRN1 levels modulate P-body formation and regulates mRNA decapping. (**A**) Western blot analysis of XRN1^KO^ HeLa cells generated with CRISPR/Cas9. (**B**) Images generated through confocal microscopy of WT and XRN1^KO^ HeLa cells. Immunofluorescent staining was performed using ⍺-EDC4 to detect P-bodies. Images were processed, and channels merged using Fiji. (**C**) Quantification of P-body sizes of WT and XRN1^KO^ HeLa cells. Measurements were performed on three separate biological replicates with ten cells per replicate using Fiji. Error bars represent the SEM and statistical significance was established using a two-tailed student’s T-test (* = P < 0.05). (**D**) Western blot analysis of WT and XRN1^KO^ HeLa cells transfected with plasmid encoding FLAG-tagged NBDY. (**E**) Images generated through confocal microscopy of WT and XRN1^KO^ HeLa cells transfected with plasmid encoding FLAG-tagged NBDY. Immunofluorescent staining was performed using ⍺-FLAG and ⍺-EDC4. Transfected cells are outlined in yellow to contrast P-body formation with non-transfected cells (blue outlines). Images were processed, and channels merged using Fiji. (**F**) mRNA decay analysis of WT and XRN1^KO^ HeLa cells transfected with plasmid encoding FLAG-tagged NBDY. RNA was isolated from cells after treatment with Actinomycin D (Act-D) to halt transcription. mRNA levels were measured by RT-qPCR with RL-6xB levels being normalized to *GAPDH* levels. Normalized RL-6xB levels at zero-hours of Act-D treatment were set to 100%. Error bars represent the SEM of three biological replicates and statistical significance was established using a two-tailed student’s T-test compared to zero hour conditions (*** = P < 0.001). (**G**) Cap status analysis of mRNAs collected from XRN1^KO^ HeLa cells transfected with control plasmids or encoding FLAG-tagged NBDY after six hours of Act-D treatment. Isolated RNA was incubated *in vitro* with terminator nuclease (T. Nuc) along with no-enzyme control samples. Purified RNA underwent RT-qPCR analysis for the annotated genes normalized to an *in vitro* transcribed *firefly luciferase* mRNA spike-in control introduced at the RT step. Normalized mRNA levels of no-enzyme control samples were set to 100%, error bars represent the SEM of three biological replicates, and statistical significance was established using a two-tailed student’s T-test relative to control (*** = P < 0.001).

### The EDC4-XRN1 interaction coordinates mRNA decapping with decay in human cells

One possible explanation for why we observe enlarged P-bodies in XRN1^KO^ cells is that increasing the stability of P-body enriched mRNAs provides P-body proteins with more mRNA to aggregate with, increasing multivalency for condensate formation. To test this hypothesis, we transfected XRN1^KO^ cells with the RL-6xB reporter, along with plasmids encoding FLAG-tagged wild-type XRN1 (F-XRN1^WT^) or a catalytically inactive XRN1 mutant (F-XRN1^E178Q^) that cannot promote mRNA decay (Chang *et al*, 2011) (**Figures 7A and S2A-B**). As anticipated, F-XRN1^WT^ rescued RL-6xB mRNA decay, whereas F-XRN1^E178Q^ failed to do so (**Figure 7B**). However, both F-XRN1^WT^ and F-XRN1^E178Q^ similarly restored P-body size (**Figures 7C and S2D**) and rescued mRNA decapping as assessed by Terminator nuclease assays (**Figure 7D**). This indicates that XRN1 does not require its exonuclease activity to regulate P-body formation and enhance mRNA decapping.

**Figure 7.**
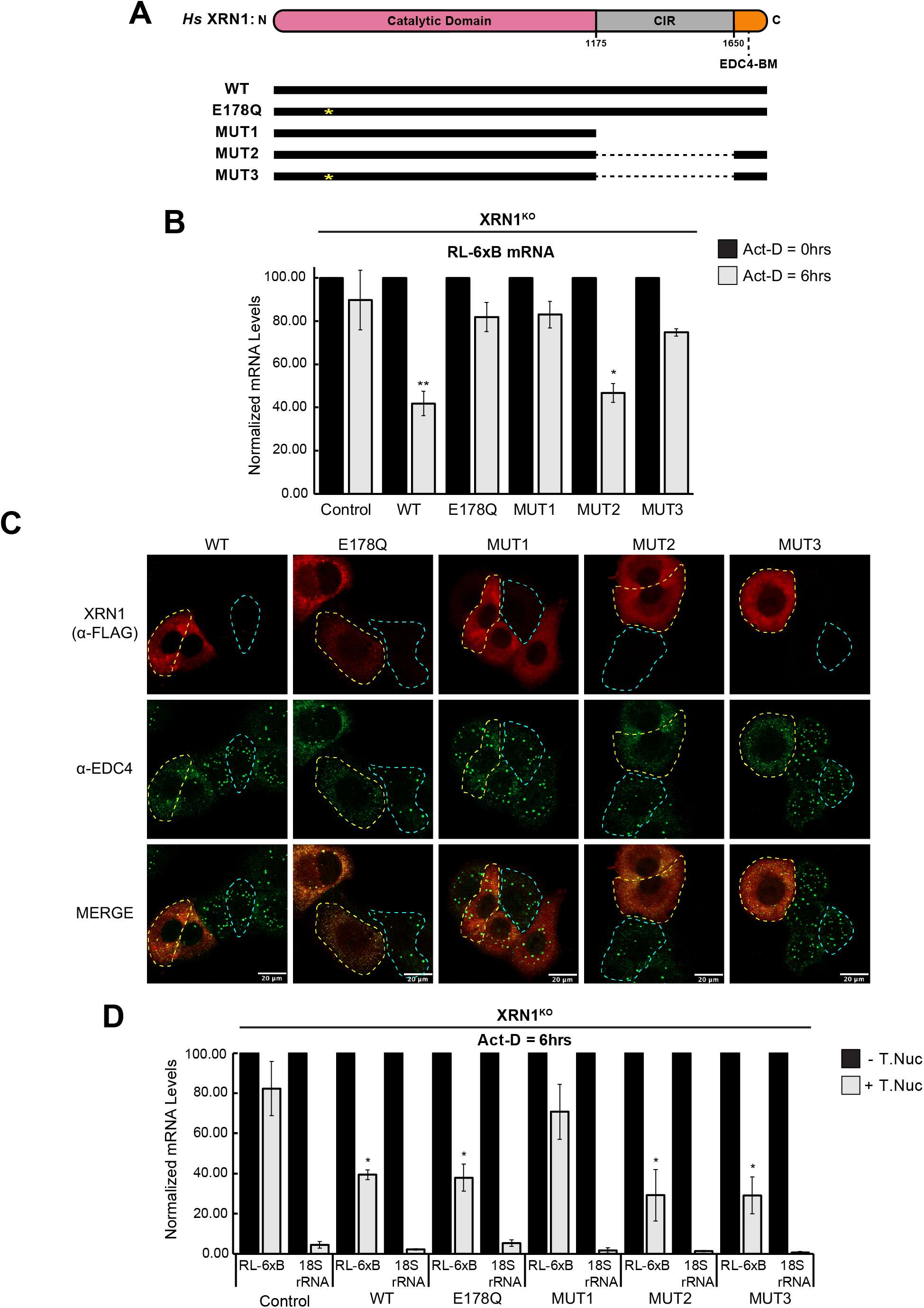
The EDC4-XRN1 interaction remodels P-bodies to regulate mRNA decapping. (**A**) Schematic diagram of human XRN1, along with diagrams representing deletion mutants expressed in the following experiments. (**B**) mRNA decay analysis of XRN1^KO^ HeLa cells transfected with plasmids encoding the annotated FLAG-tagged XRN1 proteins. RNA was isolated from cells after treatment with Actinomycin D (Act-D) to halt transcription. mRNA levels were measured by RT-qPCR with RL-6xB levels being normalized to *GAPDH* levels. Normalized RL-6xB levels at zero-hours of Act-D treatment were set to 100%. Error bars represent the SEM of three biological replicates and statistical significance was established using a two-tailed student’s T-test relative to control conditions after six hours of Act-D treatment (* = P < 0.05; ** = P < 0.01). (**C**) Images generated through confocal microscopy of XRN1^KO^ HeLa cells transfected with plasmids encoding the annotated FLAG-tagged XRN1 proteins. Immunofluorescent staining was performed using ⍺-FLAG and ⍺-EDC4. Transfected cells are outlined in yellow to contrast P-body formation with non-transfected cells (blue outlines). Images were processed, and channels merged using Fiji. (**D**) Cap status analysis of mRNAs collected from XRN1^KO^ HeLa cells transfected with plasmids encoding FLAG-tagged XRN1 proteins after 6-hours of Act-D treatment. Isolated RNA was incubated *in vitro* with terminator nuclease (T. Nuc) along with no-enzyme control samples. Purified RNA underwent RT-qPCR analysis for the annotated genes normalized to an *in vitro* transcribed *firefly luciferase* mRNA spike-in control introduced at the RT step. Normalized mRNA levels of no-enzyme control samples were set to 100%, error bars represent the SEM of three biological replicates, and statistical significance was established using a two-tailed student’s T-test relative to control samples incubated with T. Nuc (* = P < 0.05).

The Izaurralde Lab previously demonstrated that XRN1 coordinates mRNA decapping and decay in *Drosophila* by directly binding to DCP1 (Braun *et al*., 2012). In contrast, while human XRN1 also interacts with mRNA decapping factors, it does not directly interact with DCP1. Rather, XRN1 associates with the mRNA decapping complex using a conserved C-terminal EDC4-binding motif (EDC4-BM) in human cells (Braun *et al*., 2012; Chang *et al*., 2014) (**Figure 7A**). As our data show that modulating the levels of EDC4 or XRN1 regulates mRNA decapping by altering P-body formation, we next tested whether the EDC4-XRN1 interaction is responsible for regulating these processes in human cells. To this end, we complemented XRN1^KO^ cells with the XRN1 catalytic domain alone (F-XRN1^MUT1^), which lacks the EDC4-binding motif (EDC4-BM) (**Figures 7A and S2C**). However, this mutant failed to restore P-body size or promote RL-6xB mRNA decapping and decay (**Figures 7B-D and S2D**). On the other hand, fusing the XRN1 catalytic domain to the EDC4-BM (F-XRN1^MUT2^) restored P-body size and reestablished RL-6xB mRNA destabilization to levels seen with F-XRN1^WT^ rescue (**Figures 7B and C**). Importantly, a nuclease-dead version of this construct (F-XRN1^MUT3^) failed to rescue RL-6xB mRNA decay but restored both P-body size and RL-6xB mRNA decapping as assessed via Terminator nuclease assays (**Figures 7B-D and S2D**). Collectively, these data indicate that the interaction between EDC4 and XRN1 regulates P-body formation, which in turn coordinates mRNA decapping and decay in human cells.

### P-bodies support cell viability in the absence of XRN1

Stress granules are RNP granules that form in response to a variety of cellular stresses including oxidative stress, viral infection, and amino acid starvation, and contain several proteins that are also found in P-bodies (Beckham & Parker, 2008; Protter & Parker, 2016). We observed that overexpressing EDC4 or knocking out XRN1 led to an increase in P-body size. However, while overexpressing EDC4 did not lead to the formation of G3BP1-positive stress granules (**Figure S1C**), we observed a small percentage (∼5%) of XRN1^KO^ cells that contained stress granules, even though they were not exposed to cellular stressors (**Figure 8A**). One possible explanation for this is that altered P-body morphology in the absence of XRN1 can lead to low levels of cellular stress, which in turn results in stress granule formation. To test this, we expressed NBDY in XRN1^KO^ cells to disrupt P-body formation and determine whether this might attenuate stress granule formation. Incredibly, expressing NBDY in XRN1^KO^ HeLa cells led to a dramatic increase in stress granule formation, with the majority of cells displaying G3BP1-positive stress granules staining for NBDY expression (**Figures 8B and C**). In contrast, we did not observe stress granules forming in NBDY-expressing wild-type cells (**Figure 8B, top panels**). Importantly, disrupting P-body formation led to a significant decrease in viability of XRN1^KO^ cells but had no impact on wild-type cell viability (**Figure 8D**). Furthermore, these observations are specifically due to the loss of XRN1, as rescuing XRN1 levels in XRN1^KO^ cells with exogenous F-XRN1^WT^ prevented stress granules from forming (**Figure 8A**) and rescued their viability in the absence of visible P-bodies (**Figure 8D**). Thus, these data demonstrate that P-bodies support cell viability in the absence of XRN1 and prevent the formation of stress granules in the absence of external stress.

**Figure 8.**
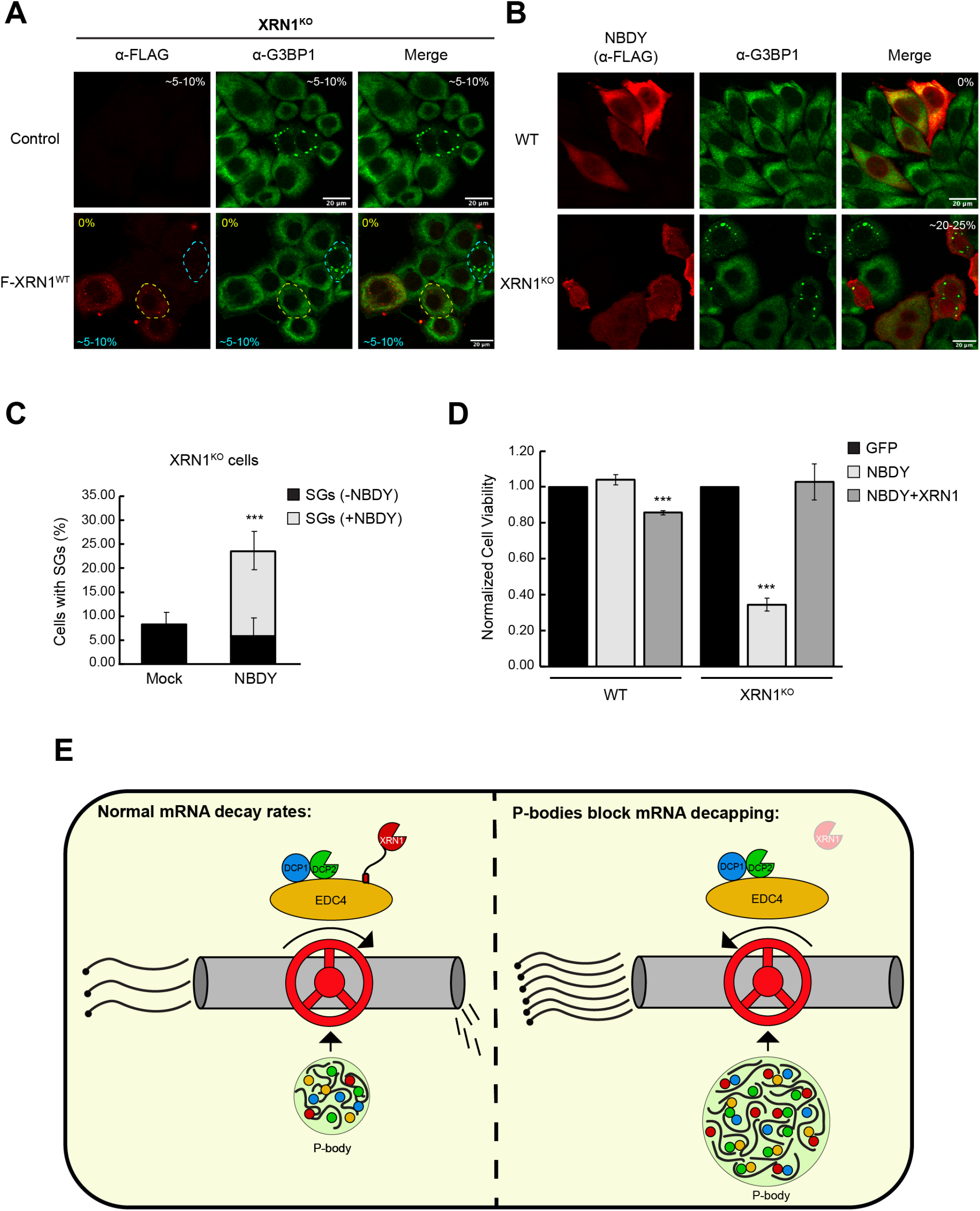
P-bodies block stress granule formation and support cellular fitness in XRN1^KO^ cells. (**A**) Images generated through confocal microscopy of XRN1^KO^ HeLa cells transfected with or without plasmids encoding the annotated FLAG-tagged XRN1^WT^. Immunofluorescent staining was performed using ⍺-FLAG and ⍺-G3BP1. Transfected cells are outlined in yellow to contrast stress granule formation (0%) with non-transfected cells (blue outlines; 5-10%). Images were processed, and channels merged using Fiji. (**B**) Images generated through confocal microscopy of WT or XRN1^KO^ HeLa cells transfected with or without plasmids encoding the annotated FLAG-tagged NBDY. Immunofluorescent staining was performed using ⍺-FLAG and ⍺-G3BP1. The percentage of cells displaying stress granules is annotated under each condition. Images were processed, and channels merged using Fiji. (**C**) Quantification of the percentage of cells that form stress granules in XRN1^KO^ cells under mock transfection and NBDY transfection conditions. Black bars indicate the percentage of cells with stress granules that occurs under control conditions (left) or negative for FLAG-NBDY staining (right), with the gray bar indicating the percentage of cells with SGs that stained positive for FLAG-NBDY expression. Error bars represent the SEM of three biological replicates and statistical significance was determines using a two-tailed student’s T-test relative to mock transfection (*** = P < 0.001). (**D**) Fluorometric cell viability analysis of WT and XRN1^KO^ cells transfected with plasmids encoding the indicated proteins. Cell viability was normalized to 1.0 for cells expressing GFP-control plasmids. Error bars represent the SEM of three biological replicates and statistical significance was determines using a two-tailed student’s T-test relative to GFP-control cells (*** = P < 0.001). (**E**) Model representing how P-bodies respond to the loss of XRN1 to regulate mRNA decapping and promote cellular fitness.

## DISCUSSION

Descriptions of a mechanistic role(s) for P-bodies in regulating mRNA stability has been historically elusive. In this study we present data that support a model whereby P-bodies regulate mRNA decapping in response to changes in mRNA decapping and decay factor stoichiometry (**Figure 8E**). We show that enhancing P-body formation by overexpressing EDC4, depleting XRN1, or disrupting the EDC4-XRN1 interaction impairs mRNA decapping of P-body-enriched mRNAs. Together, our data point to a model whereby EDC4-XRN1 stoichiometry and their association feeds back on mRNA decapping activity by modulating P-body morphology. Moreover, our data suggest that enhanced P-body formation in the absence of XRN1 attenuates stress granule formation and promotes cellular fitness. To our knowledge, this study is the first to directly attribute a mechanistic function to P-bodies in the post-transcriptional regulation of gene expression.

Transcriptomic analyses of P-body enriched mRNAs demonstrate that they are mainly comprised of poorly translated mRNAs that are targeted for decay by RNA-binding proteins via *cis*-regulatory elements (Hubstenberger *et al*., 2017; Matheny *et al*, 2019). However, these analyses suggest that mRNAs within P-bodies are not actively undergoing decay. Importantly, sequencing read distribution of transcripts purified directly from P-bodies was significantly shifted toward the 5’end of the mRNAs relative to the total and P-body-depleted transcriptomes, suggesting that they are being protected from 5’ to 3’ decay (Hubstenberger *et al*., 2017). In line with these observations our results show that P-bodies can be remodeled to impair mRNA decapping, leading to further mRNA enrichment within P-bodies that protects them from 5’ to 3’ decay. However, precisely how P-bodies block mRNA decapping remains unknown. *In vitro* studies using recombinant yeast DCP1 and DCP2 proteins suggest that liquid-liquid phase separation can bias DCP2 conformation to impair its pyrophosphatase activity (Tibble *et al*., 2021). P granules in *C. elegans* germ cells—structures compositionally similar to P-bodies—have been shown to exhibit liquid like behaviors that rapidly dissolve and condense (Brangwynne *et al*, 2009). Therefore, P-body phase transitions may alter DCP2 conformation or localization *in vivo*, which could explain how P-body aggregation inhibits mRNA decapping. Another plausible explanation is that P-bodies serve a physical barrier that limits DCP2 from interfacing with its target mRNAs. In support of the latter hypothesis, mRNAs targeted by DCP2 are disproportionately enriched within P-bodies (Luo *et al*, 2020). Moreover, a large proportion of DCP2-target mRNAs are also stabilized by knocking out NBDY, which leads to increases in the number and sizes of P-bodies (Na *et al*., 2020). Therefore, enhanced P-body enrichment of DCP2 target mRNAs may be what prevents their decapping. Nevertheless, additional research is needed to determine the structural underpinnings of mRNA decapping regulation by P-bodies *in vivo*.

We have shown that the EDC4-XRN1 interaction is a key regulator of P-body size and mRNA decapping activity. XRN1^KO^ human cells exhibit significantly larger P-bodies and deficits in decapping of miRNA-targeted mRNAs, both of which can be rescued by XRN1 variants that re-establish contact with EDC4—irrespective of XRN1 exoribonuclease activity. These results are in line with studies from the Izarraulde Lab, which showed that an XRN1 mutant that cannot bind EDC4 failed to degrade a reporter mRNA targeted by the nonsense mRNA decay protein SMG7 in human cells (Chang *et al*., 2019). Though they did not assess whether this XRN1 mutant failed to degrade the SMG7-targeted reporter was due to deficits in mRNA decapping activity, previous work demonstrated that the DCP1-XRN1 interaction is required for efficient mRNA decapping in *Drosophila* S2 cells (Braun *et al*., 2012). However, this interaction is not conserved in vertebrates, where XRN1 instead directly binds EDC4 (Braun *et al*., 2012; Chang *et al*., 2014). Interestingly, XRN1 binds to the C-terminal ⍺-helical domain of EDC4, a region that has been reported to oligomerize with other EDC4 proteins and is required for P-body formation (Brothers *et al*., 2022; Chang *et al*., 2014). In keeping with what we observed upon XRN1 depletion, our results indicate that overexpressing EDC4 leads to larger P-bodies that inhibit decapping activity. This suggests that reducing the amount of XRN1 bound to EDC4—either by depleting XRN1 or by increasing EDC4 levels—is what dictates P-body size and decapping activity, potentially by increasing the rate of EDC4 oligomerization. This raises the possibility that EDC4 may respond to changes in XRN1 levels by enhancing P-body formation as a feedback mechanism to block untimely or unwanted mRNA decay. This could occur under conditions where the decapping complex subunit levels change dramatically, or when XRN1 levels simply cannot keep up with the number of mRNAs that are targeted for decay. However, more research is needed to provide additional support for such a model.

Despite observations that enhancing P-body formation stabilizes miRNA-targeted mRNAs by blocking decapping, we did not observe changes in the luciferase outputs produced by RL-6xB reporter mRNAs when manipulating P-body formation. Rather, our data suggest that P-body remodeling shifts the mode of miRNA-silencing from decay to translational repression. This is consistent with observations that P-body enriched mRNAs tend to be poorly translated and that P-bodies do not contain ribosomal proteins (Hubstenberger *et al*., 2017; Matheny *et al*., 2019; Youn *et al*, 2018). Indeed, single-molecule imaging studies of reporter mRNAs tethered by the miRNA machinery suggest that P-bodies are sites of persistent translational repression for miRNA-targeted mRNAs (Cialek *et al*., 2022). Notably, the first step in miRNA-mediated mRNA decay involves the recruitment of deadenylases, with deadenylation being associated with translational repression in mammalian cell free extracts and *Xenopus* oocytes (Chekulaeva *et al*, 2011; Cooke *et al*, 2010; Fabian *et al*., 2009). In agreement with these observations, our results suggest that miRNA-targeted mRNAs stabilized by P-bodies are deadenylated, which may coincide with them being translationally repressed. In addition to miRNA-targeted mRNAs, mRNAs bound by the core P-body protein 4E-T are also translationally repressed in a deadenylated state (Rasch *et al*, 2020). Consistent with our data, 4E-T-bound mRNAs remained capped, dependent on the ability of 4E-T to bind and recruit the cap binding proteins eIF4E and 4EHP to P-bodies. Several reports across a diverse range of biological contexts suggest that disassembling P-bodies in response to cellular or developmental cues may allow translationally repressed P-body mRNAs to re-initiate their translation (Buddika *et al*, 2021; Di Stefano *et al*., 2019; Sankaranarayanan *et al*, 2021). Thus, it is possible that P-bodies inhibit decapping and recruit the cap-binding protein eIF4E so that P-body enriched mRNAs can undergo a pioneering round of translation before being decayed in biological or developmental contexts where P-bodies dissolve. However, this hypothesis has not been directly tested.

While disrupting P-body formation has not been previously linked to impacting cell viability, our data here point to P-bodies supporting cell fitness. Expressing NBDY in XRN1^KO^ cells not only disrupted P-bodies but also led to the formation of stress granules and a concomitant decrease in cell viability. Interestingly, while we were able to generate XRN1^KO^ or LSM14A^KO^ cells lines (the latter of which fails to form visible P-bodies), we were unable to generate homozygous XRN1/LSM14 double knockouts in either genetic background (data not shown). This suggests that loss of both XRN1 and P-body formation may confer a synthetic lethal phenotype. Preventing P-bodies from forming may lead to a build-up of stable decapped mRNAs in cells lacking XRN1. As these transcripts would theoretically be unable to initiate translation in the absence of their 5’-caps, it is possible that they could act to nucleate stress granules in the absence of stress. Nevertheless, exactly how P-bodies buffer stress granule formation and support cell viability in the absence of XRN1 remains to be elucidated.

In conclusion, our results show that P-bodies dynamically regulate mRNA decapping in response to changes in decapping complex stoichiometry and association. Future studies are needed to establish to what degree the regulation of decapping is involved in the post-transcriptional regulation of gene expression in biological contexts where cytoplasmic granule remodeling occurs. Moreover, our work suggests that P-bodies buffer stress granule formation and support cellular fitness in the absence of XRN1. Future research will be needed to understand how P-bodies can compensate for disrupted 5’ to 3’ decay to support cellular viability.

## METHODS

### Cell Lines and Cell Culture

We obtained epithelioid carcinoma HeLa cells from ATCC. Cell lines were identified via morphology but have not been authenticated. Cells were assessed and tested negative for mycoplasma contamination. HeLa cells were grown in Dulbecco’s modified Eagle’s medium (DMEM) supplemented with 10% fetal bovine serum, 50 U/mL penicillin, and 50 mg/mL streptomycin. FLAG-LSM14A complementation cell lines were generated as previously described (Brandmann *et al*., 2018).

### Plasmids and Antibodies

All plasmid transfections were performed using polyethyleneimine (PEI). V5-tagged EDC4 encoding plasmids were generated by Gateway cloning from pDONR-EDC4 into a pDEST40 vector. FLAG-tagged XRN1 constructs were generated by cloning PCR amplicons into pQCXIB using the AgeI and EcoRI sites. FLAG-tagged NBDY was cloned into pcDNA3.1 as previously described (Brothers *et al*., 2022). RL-6xB, RL-6xBMUT, and FL reporter plasmids were constructed as previously described (Fabian *et al*., 2009; Pillai *et al*, 2005). Antibodies for this study were purchased against FLAG (Sigma), V5 (Rabbit – Cell Signaling; Mouse – Thermofisher), EDC4 (Bethyl), DDX6 (Bethyl), XRN1 (Bethyl), LSM14 (GeneTex), ACTIN (Sigma), and GAPDH (Cell Signaling). Antibodies used in western blot and immunofluorescence microscopy were diluted to the manufacturer’s specifications.

### CRISPR-Cas9 Gene Editing

gRNAs targeting LSM14A (5’-TCTGTACCAAAGGATCGAAC-3’) and XRN1 (5’-AGAGAAGAAGTTCGATTTGG-3’) were cloned into LentiCRISPR v2 (Addgene #52961) using BsmBI restriction enzyme sites and confirmed by sequencing. WT HeLa cells were transiently transfected with recombinant LentiCRISPR v2 plasmids followed by puromycin (2µg/mL) selection 48 hours after transfection. 48 hours after selection, cells were returned to normal cell culture medium to recover for 24 additional hours. After selection recovery, cell lines were plated cells into 96-well plates, with monoclonal lines being grown up and screened by western blotting for knockouts.

### Immunofluorescent and smFISH microscopy

Standard immunofluorescent (IF) analyses were performed as previously described (Brothers *et al*., 2022). Concomitant IF and smFISH analyses were adapted from (Adivarahan *et al*, 2018): Previously transfected HeLa cells were plated on coverslips and grown for 12-hours to allow them to adhere. Coverslips were washed with 1xPBS and fixed with 4% PFA/1xPBS for 10min at RT. Coverslips were washed with 1xPBS and then stored in 1mL of 70% EtOH overnight at −20℃. EtOH was removed and cells were washed with 1xPBS before being air dried for 5min. This was followed by permeabilization with 0.5% Triton x-100/1× PBS for 10 min at RT and three washes of 1xPBS. Hybridization buffers were prepared as previously described (Adivarahan *et al*., 2018) using Cy3-labeled probes complimentary to the *Renilla* luciferase CDS (Biosearch/Stellaris). Coverslips were incubated with hybridization buffer for 3 hours at 37℃ shielded from light. Coverslips were then washed twice with 10% Formamide/2xSSC for 30min per wash at 37℃ shielded from light, followed by three washes with 1xPBS (no incubation). Hybridized coverslips were then blocked with 4% BSA for 10min at RT followed by an overnight incubation at 4℃ with primary antibodies diluted in 1% BSA (shielded from light). After primary antibody incubation, coverslips were washed with 1xPBS and incubated with Alexa Fluor 488 goat anti-rabbit and Alexa Fluor 594 goat anti-mouse antibodies diluted 1:500 in 1% BSA for 45min at RT (shielded from light). This was followed by washing the coverslips with 1xPBS, and nuclei were stained with DAPI for 15 min at RT (shielded from light). Coverslips were washed with 1xPBS before being mounted onto glass slides with ProLong Gold media (ThermoFisher). Images were taken using a Zeiss Confocal LSM 800 microscope at 40X magnification and processed with Fiji to merge channels, add scale bars, and quantify smFISH RL-6xB signal.

### Luciferase Assays and mRNA Decay Assays

HeLa cells were transfected with the indicated plasmids at 30% confluency. 24 hours after transfection, cells were reseeded into 6-well dishes for collection at the relevant timepoints in mRNA decay assays, along with additional wells for luciferase assays. 24 hours later, cells were harvested and lysed in Passive Lysis Buffer (Promega). The activity levels of the *Renilla* (RL) and firefly (FL) luciferase was measured using a Dual-Luciferase Assay (Promega). For mRNA decay assays, cell culture media was replaced with media containing Actinomycin D (5 µg/mL). Cells were harvested at the indicated timepoints, pelleted, and flash frozen before being stored at - 80°C. Frozen pellets were processed for RT-qPCR analyses as previously described (Brothers *et al*., 2022).

### Poly(A) Tail Analyses

Enhanced poly(A) tail (ePAT) analyses were performed as previously described (Jänicke *et al*., 2012). The anchor primer used in the ePAT reverse transcription step is (5’- GCGAGCTGGCGCCGGCGCTTTTTTTTTTTT-3’). The (dT)12VN primer used in the reverse transcription to generate a size marker for the fixed length A_12_-tail product is (5’- GCGAGCTGGCGCCGGCGCTTTTTTTTTTTTVN-3’). PCR reactions were then performed using Taq polymerase (Biobasic), resolved on 2% agarose gels stained with EtBr. Gels were visualized using an ImageQuant LAS 4000 imager (GE Healthcare).

### Terminator Nuclease Assays

Purified RNA was incubated with Terminator nuclease and buffer A (Biosearch Technologies) for 1hr at 30℃, according to the manufacturer’s specifications. RNA was then resuspended in an equal volume of 3M NaOAc (pH 5.5) and phenol chloroform extracted. Purified RNA was EtOH precipitated and resuspended in water before undergoing reverse transcription (RT) with an *in vitro* transcribed *firefly luciferase* mRNA spike-in control (0.1ng per reaction) to normalize for RT efficiency. Quantitative PCR (qPCR) was subsequently performed as described above, except for 18S rRNA analysis where cDNA was diluted 1:1000 before undergoing qPCR. For recombinant mRNA decapping enzyme incubation, purified RNA was incubated with mRNA decapping enzyme and buffer (NEB) for 1hr at 37℃, according to the manufacturer’s specifications. RNA was then resuspended in an equal volume of 3M NaOAc (pH 5.5) and phenol chloroform extracted. Purified RNA was EtOH precipitated and resuspended in water before being divided and incubated with Terminator nuclease, as described above.

### Fluorometric Cell Viability Assays

Cells were carefully counted and seeded at equal density. 24-hours after seeding, cells were transfected with plasmids encoding the indicated proteins along with a puromycin selection cassette. 24-hours post-transfection, transfected cells were selected for with media containing puromycin (2 µg/mL) and grown for an additional 48-hour. Cells were then harvested, stained with acridine orange and propidium iodide and live cells were counted CellDrop FL Cell Counter (Denovix).

---

**Supplemental Figure 1.**
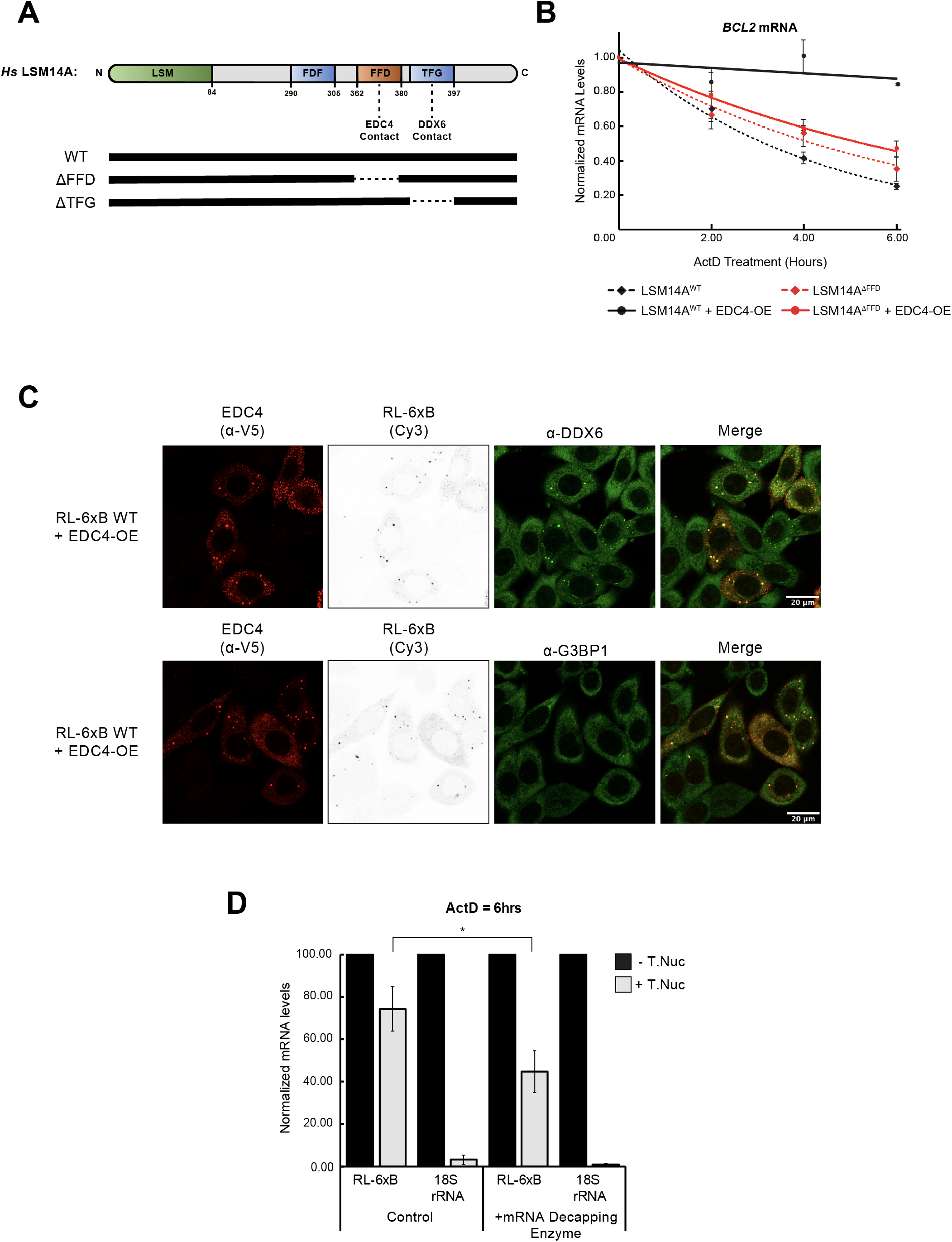
(**A**) Duplication of the schematic diagram of human LSM14A (**Figure 2A**), along with diagrams representing deletion mutants that disrupt P-body formation. (**B**) mRNA decay assays in the indicated F-LSM14 cell lines assessing the decay rates of *BCL2* mRNAs with or without EDC4-overexpression (EDC4-OE). RNA was isolated from cells after treatment with Actinomycin D (Act-D) to halt transcription. mRNA levels were measured by RT-qPCR with *BCL2* levels being normalized to *GAPDH* levels. Normalized *BCL2* levels at zero-hours of Act-D treatment were set to 1.0. Error bars represent the SEM of three biological replicates. (**C**) Images generated through confocal microscopy of F-LSM14A^WT^ HeLa cells transfected with plasmids encoding EDC4-V5. Single-molecule fluorescent *in situ* hybridization (smFISH) using Cy3-labeled probes against RL-6xB with simultaneous immunofluorescent staining was performed using ⍺-V5 antibodies with ⍺-DDX6 or ⍺-G3BP1 to differentiate between P-bodies and stress granules. Images were processed, analyzed, and channels merged using Fiji. (**D**) Cap status analysis of mRNAs collected from HeLa cells overexpressing EDC4 after 6-hours of Act-D treatment. Isolated RNA was incubated *in vitro* with mRNA decapping enzyme followed by terminator nuclease (T. Nuc) along with no-enzyme controls for both incubations. Purified RNA underwent RT-qPCR analysis for the annotated genes normalized to an *in vitro* transcribed *firefly luciferase* mRNA spike-in control introduced at the RT step. Normalized mRNA levels of no T. Nuc control samples were set to 1.0, error bars represent the SEM of three biological replicates, and statistical significance was established using a two-tailed student’s T-test (* = P < 0.05).

**Supplemental Figure 2.**
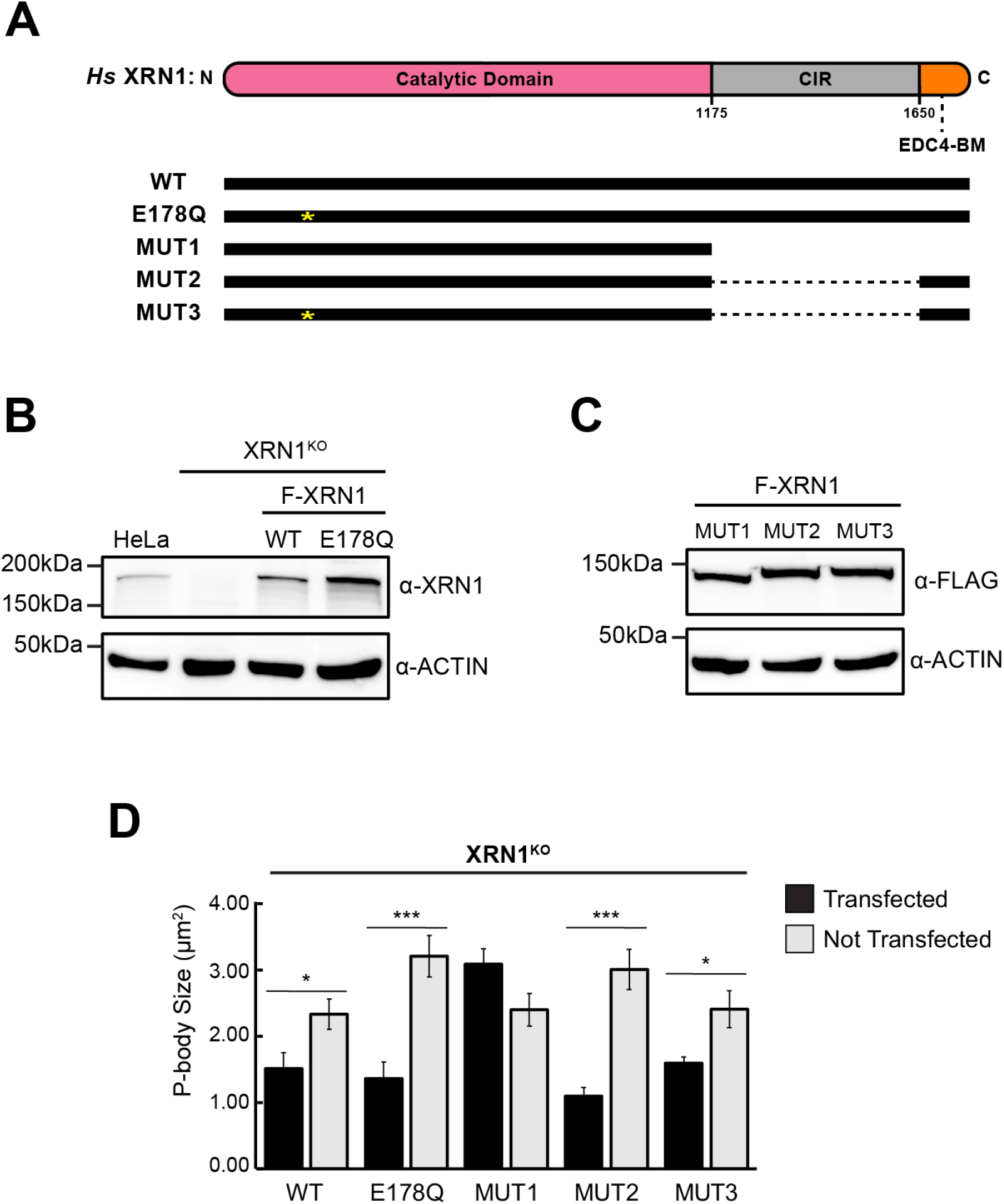
(**A**) Duplication of the schematic diagram of human XRN1 (**Figure 7A**), along with diagrams representing deletion mutants expressed in the following experiments. (**B**) Western blot analysis of WT or XRN1^KO^ cells expressing full-length F-XRN1 variants. (**C**) Western blot analysis of truncated F-XRN1 mutants. (**D**) Quantification of P-body sizes of XRN1^KO^ HeLa cells expressing the indicated F-XRN1 proteins. Measurements were performed on three separate biological replicates with ten cells per replicate using Fiji. Error bars represent the SEM and statistical significance was established using a two-tailed student’s T-test relative to matched non-transfected control cells in the same field of view (* = P < 0.05).

